# Single-cell transcriptomics reveals a mechanosensitive injury signaling pathway in early diabetic nephropathy

**DOI:** 10.1101/2022.10.19.512894

**Authors:** Shuya Liu, Yu Zhao, Shun Lu, Tianran Zhang, Maja T. Lindenmeyer, Viji Nair, Sydney E. Gies, Guochao Wu, Robert G. Nelson, Jan Czogalla, Hande Aypek, Stephanie Zielinski, Zhouning Liao, Melanie Schaper, Damian Fermin, Clemens D. Cohen, Denis Delic, Christian F. Krebs, Florian Grahammer, Thorsten Wiech, Matthias Kretzler, Catherine Meyer-Schwesinger, Stefan Bonn, Tobias B. Huber

## Abstract

Diabetic nephropathy (DN) is the leading cause of end-stage renal disease and histopathologic glomerular lesions are among the earliest structural alterations of DN. However, the signaling pathways that initiate these glomerular alterations are incompletely understood. To delineate the cellular and molecular basis for DN initiation, we performed single-cell and bulk RNA sequencing of renal cells from type 2 diabetes mice (BTBR *ob/ob*) at the early stage of DN. Analysis of differentially expressed genes revealed glucose-independent responses in glomerular cell types. The gene regulatory network upstream of glomerular cell programs suggested the activation of mechanosensitive transcriptional pathway MRTF-SRF predominantly taking place in mesangial cells. Importantly, activation of MRTF-SRF transcriptional pathway was also identified in DN glomeruli in independent patient cohort datasets. Furthermore, ex vivo kidney perfusion suggested that the regulation of MRTF-SRF is a common mechanism in response to glomerular hyperfiltration. Overall, our study presents a comprehensive single-cell transcriptomic landscape of early DN, highlighting mechanosensitive signaling pathways as novel targets of diabetic glomerulopathy.

## Introduction

Diabetic nephropathy (DN) is one of the major complications in diabetic patients and is the leading cause of end-stage renal disease worldwide [1]. DN is a complex disease and its progression, particularly in type 2 diabetes, is confounded by multiple pathogenic factors, such as insulin resistance and obesity [1, 2]. Previous studies used single-cell or single-nucleus RNA sequencing (scRNA-seq or snRNA-seq) to investigate cellular changes or drug responses in diabetic kidneys. Single-nucleus transcriptomics on early human diabetic kidneys suggests increased potassium secretion in the distal nephron and pro-angiogenic signaling in diverse kidney cells [3]. A following study combining single-nucleus RNA and assay for transposase-accessible chromatin (ATAC) sequencing highlights the glucocorticoid signaling in the proximal tubule [4]. Stefansson et al. provide molecular programs associated with glomerular hyperfiltration in human DN [5], and Fu et al. show the single-cell transcriptomics of eNOS -/- mice with streptozotocin-induced diabetes, which is a type 1 diabetes model without obesity nor insulin-resistance [6]. Single-cell/nucleus transcriptomics on drug treated murine diabetic kidneys suggest important alterations in proximal tubule cells [7, 8].

DN is characterized by distinct histopathological lesions dominating the renal glomerulus, such as glomerular basement membrane (GBM) thickening, mesangial expansion and nodular glomerulosclerosis [2]. Early changes such as glomerular hyperfiltration have been proposed to be critical for the subsequent development of glomerulosclerosis [1]. Preservation of glomerular filtration rate is of great importance for renoprotection [2]. However, the signaling pathways that initiate these glomerular alterations are incompletely understood. Therefore, it is particularly warranted to understand the cellular changes and molecular mechanisms that initiate DN in the glomerulus.

In this study, we conducted a comprehensive analysis of cellular changes in type 2 diabetic glomeruli. The type 2 diabetes mouse model BTBR *ob/ob* exhibits insulin resistance, hyperglycemia and obesity, as well as the rapid progression of diabetic glomerulopathy [9–11]. We generated single-cell transcriptomics of kidneys and bulk transcriptomics of purified glomerular cell types. We compared mouse DN with human DN and validated our findings in different patient cohort datasets at both mRNA and protein levels. Our study reveals the upregulation of mechanosensitive signaling pathway MRTF-SRF in response to glomerular hyperfiltration, which is associated with glomerulopathy in early DN.

## Results

### scRNA-seq of BTBR ob/ob mouse kidneys with early DN

We generated BTBR *ob/ob* podocyte-reporter mice (BTBR.Cg-Lep^ob/ob WiscJ^;Gt (ROSA)26Sor^tm4(ACTB-tdTomato,-EGFP)Luo^;Tg(NPHS2-cre)^295Lbh^; hereafter referred to as BTBR *ob/ob* or DN mice) to investigate the cellular and molecular changes that occur in DN. By 6 weeks of age, both male and female BTBR *ob/ob* mice exhibited obesity, hyperglycemia, and albuminuria (Supplementary Fig. 1a). From 12 weeks of age, obvious glomerular hypertrophy was observed in histological analysis (Supplementary Fig. 1b). Mesangial expansion with accumulation of collagen IV was detected in the DN glomeruli (Supplementary Fig. 1c and d). Based on these observations, we defined 6 weeks of age as the onset stage of DN and 12 weeks of age as the early stage of DN.

Kidney tissue was sampled from a total of 16 mice, including male and female each of BTBR *ob/ob* (DN) and BTBR WT (control) mice at 6 and 12 weeks of age. The samples were partially depleted of the proximal tubule (PT) segment using a Percoll gradient [12] to enrich glomeruli. Single-cell suspensions were prepared using cold-active protease (CAP) to reduce dissociation-induced artifacts in kidney cells [13]. A total of 70,944 single cells were profiled after data pre-processing and quality control (Fig. 1b). Unsupervised clustering identified 21 clusters (Supplementary Fig. 2a, b and c; Methods). The cells in the 21 clusters were classified into 19 cell types and annotated on the basis of cell-specific marker genes reported in previous kidney and glomerulus single-cell data (14-16). Defining marker genes for each cell type and unbiased marker genes for each cluster are demonstrated in a dot plot (Fig. 1c) and in a heatmap (Supplementary Fig. 2d), respectively.

**Fig. 1:**
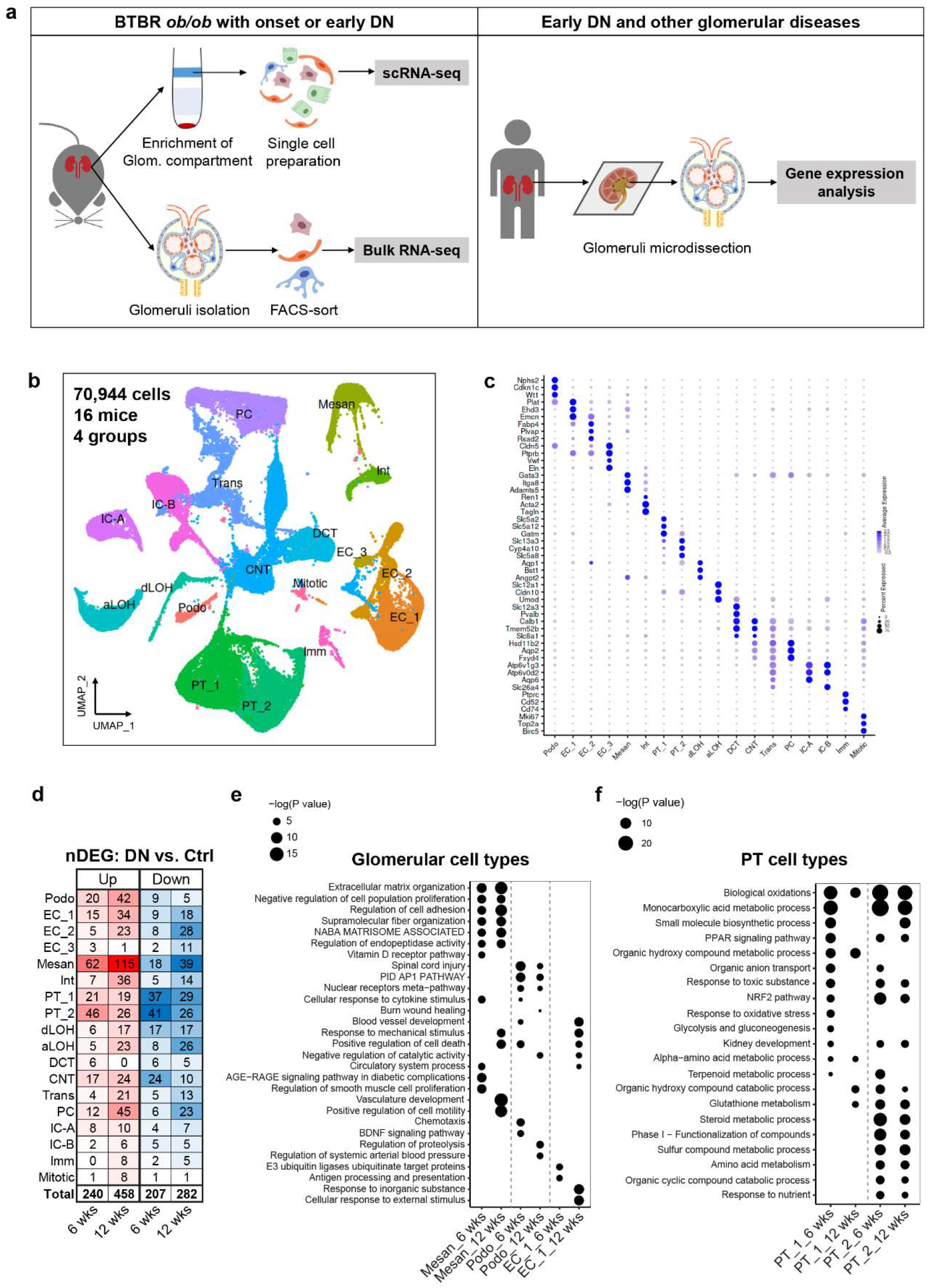
scRNA-seq of BTBR ob/ob mouse kidneys with early DN. (a) Experimental scheme. (**b**) UMAP plot of annotated cell types. (**c**) Dot plot of defining marker genes for each cell type. (**d**) Total number of significant DEGs (nDEGs) in each cell type. Top enriched pathways in glomerular (**e**) and PT cell types (**f**) at 6 and 12 weeks. UMAP: uniform manifold approximation and projection, Ctrl: control, DN: diabetic nephropathy, Podo: podocyte, EC: endothelial cell, Mesan: mesangial cell, Int: interstitial cell, PT: proximal tubule, dLOH: descending limb of loop of Henle, aLOH: ascending limb of loop of Henle, DCT: distal convoluted tubule, CNT: connecting tubule, PC: collecting duct principal cell, IC-A: A-type collecting duct intercalated cell, IC-B: B-type collecting duct intercalated cell, Trans: transition cell, Imm: immune cell, Mitotic: mitotic cell.

We identified all major components in the kidneys. The endothelial cell (EC) subtypes expressed distinct marker genes, such as *Ehd3* for EC_1 (glomerular ECs/gECs), *Plvap* for EC_2 (fenestrated ECs with diaphragm in veins and peritubular capillaries) and *Fbln2* for EC_3 (arteriolar and arterial ECs) [14]. Mesangial cells defined by marker genes *Gata3, Itga8* and *Adamts5* are clearly distinguished from interstitial cells, which were compose of vascular smooth muscle cells marked by *Acta2* and *Tagln*, and renin cells marked by *Ren1* [15, 16]. The subtype PT_1 and PT_2 were composed of cells from S1/2 and S2/3 PT segments, respectively, as they showed a continuum of marker gene expression consistent with known S1 (*Slc5a2, Slc5a12* and *Gatm*) to S3 (*Slc13a3, Cyp4a10* and *Slc5a8*) PT segments.

Due to PT depletion, the percentage of PT cells was reduced to 22.4%, while glomerular cell types were enriched to 15.6% (Supplementary Fig. 2e). The fractions of control and DN cells in each cell type were similar except for podocytes, which included more control (approximately 77%, 807 cells) than DN cells (approximately 23%, 235 cells) (Supplementary Fig. 2f). All cell types were identified in individual kidney samples (Supplementary Fig. 3a), and cells from individual kidney samples were also identified for each cell type (Supplementary Fig. 3b). The overall low expression of dissociation-induced stress genes in individual cell types (Supplementary Fig. 3c) indicates the advantages of our modified single-cell preparation protocol (Methods), which shortened the 37°C digestion time and omitted the steps stressful to glomerular cell dissociation.

### Cell type-specific alterations in DN kidneys suggest glucose-dependent and glucose-independent responses in diabetes

Next, we compared gene expression levels in DN versus control kidneys by cell type. In total, we identified 447 and 740 differentially expressed genes (DEGs) across all cell types at 6 and 12 weeks, respectively (Fig. 1d). Among all cell types, mesangial cells and PT cells had the highest numbers of DEGs (Fig. 1d), indicating that these two cell types were predominantly affected in the early stage of DN. Enrichment analysis identified significant pathways in glomerular cell types and PT cell types at 6 and 12 weeks (Fig. 1e and f). Glomerular cell types showed regulations of extracellular matrix molecules, cell adhesion, cell proliferation etc. and cellular responses to mechanic stress, cytokines and blood pressure. PT cell types showed multiple metabolic processes such as biological oxidations, and responses involved in oxidative stress, such as NRF2 and PPAR signaling pathways. These differences suggest glucose-dependent and glucose-independent responses in PT and glomerular cell types in early DN, respectively.

### Shared features of human and experimental DN

Early changes such as glomerular hypertrophy, GBM thickening and mesangial expansion occur in both human and animal DN [17]. Thus, we conducted intensive analyses on glomerular cell types.

We isolated glomeruli from BTBR *ob/ob* and BTBR WT kidneys and subsequently purified podocytes, mesangial cells and gECs by fluorescence-activated cell sorting (FACS) (Supplementary Fig. 4, Supplementary Methods). Bulk RNA-seq was performed for each purified glomerular cell type. The similarity and purity of individual glomerular cell type samples were controlled by principal component analysis (PCA) and the expression levels of specific marker genes (Supplementary Fig. 5). The cell gene expression data from scRNA-seq were compared to the bulk RNA-seq data. Podocytes, EC_1/gECs and mesangial cells showed positive Pearson correlation coefficients, suggesting a linear correlation of the two datasets (Fig. 2a). Overlapping the glomerular DEGs from single-cell and bulk RNA-seq datasets resulted in 194 unique genes (Fig. 2b), which were mapped to the European Renal cDNA Bank (ERCB) patient dataset. The mRNA expression of 106 genes was significantly regulated in microdissected glomeruli from DN patients (Fig. 2b and c). Importantly, most DEGs detected in human DN glomeruli were significantly changed in mesangial cells in both single-cell and bulk RNA-seq data (Fig. 2c).

**Fig. 2:**
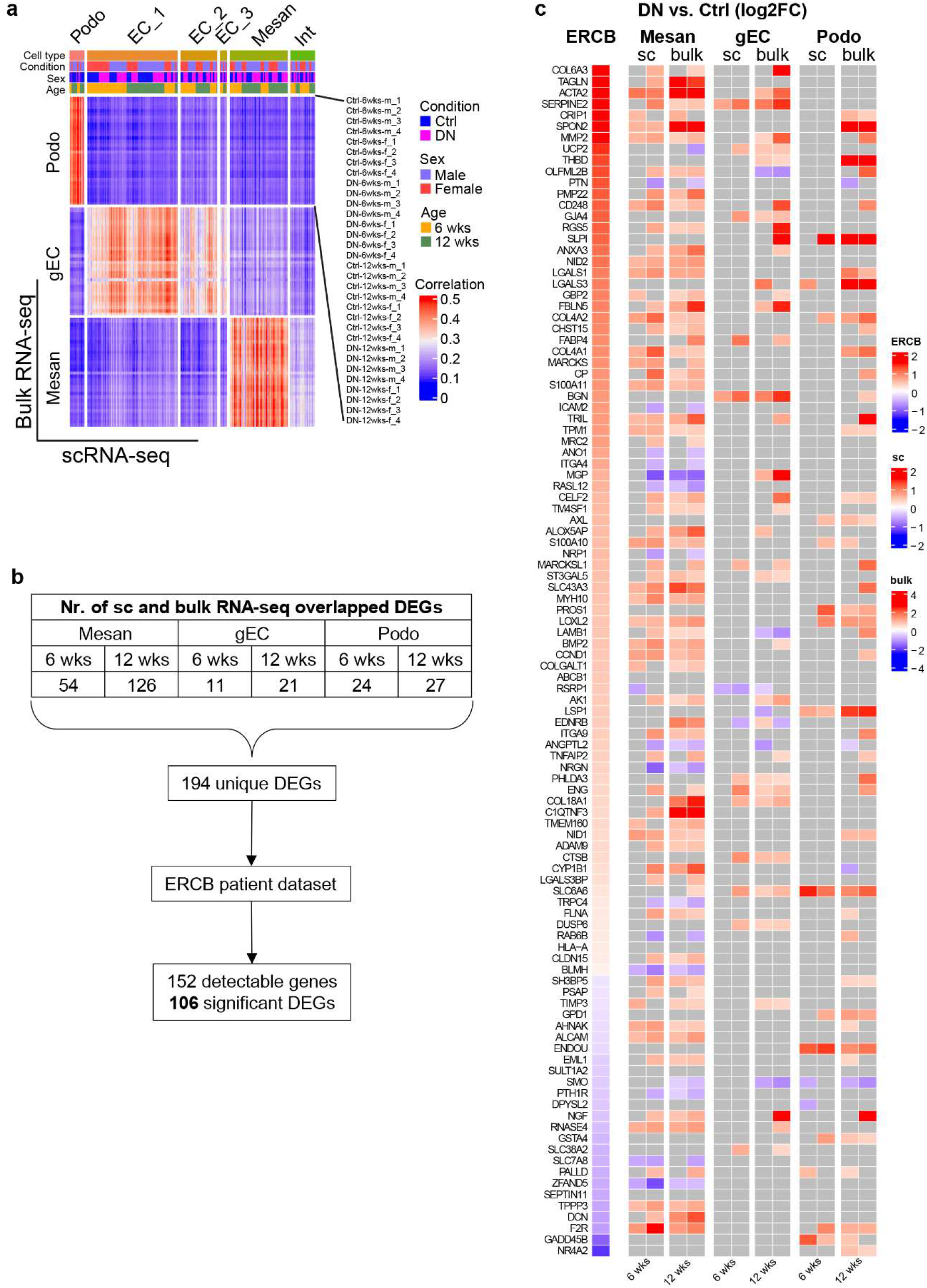
Shared features of human and experimental DN. (**a**) Heatmap displaying the Pearson correlation coefficient between scRNA-seq and bulk RNA-seq data. (**b**) Overview of significant glomerular DEGs detected in both single-cell and bulk RNA-seq datasets, as well as in the ERCB patient dataset. (**c**) Heatmap showing the significant regulation of DEGs identified in microdissected glomeruli and in glomerular cells from DN patients and mice, respectively. Nonsignificant genes are shown in grey. ERCB: European Renal cDNA Bank.

### Mechanosensitive transcriptional regulators are activated in DN glomeruli

To identify the transcriptional regulations responsible for the differential expression of genes in DN glomerular cell types, we performed upstream analysis using Ingenuity Pathway Analysis (IPA), which identifies not only transcription factors but also transcription coregulators such as coactivators. Transcriptional regulators commonly changed in single-cell and bulk RNA-seq data are shown in Figure 3a. Changes in transcriptional regulations were predominant in mesangial cells. Intriguingly, we observed the activation of mechanosensitive transcriptional regulators, including coactivators myocardin-related transcription factor A and B (MRTFA/B) and yes-associated protein 1 (YAP1 or YAP), as well as the transcription factor serum response factor (SRF).

**Fig. 3:**
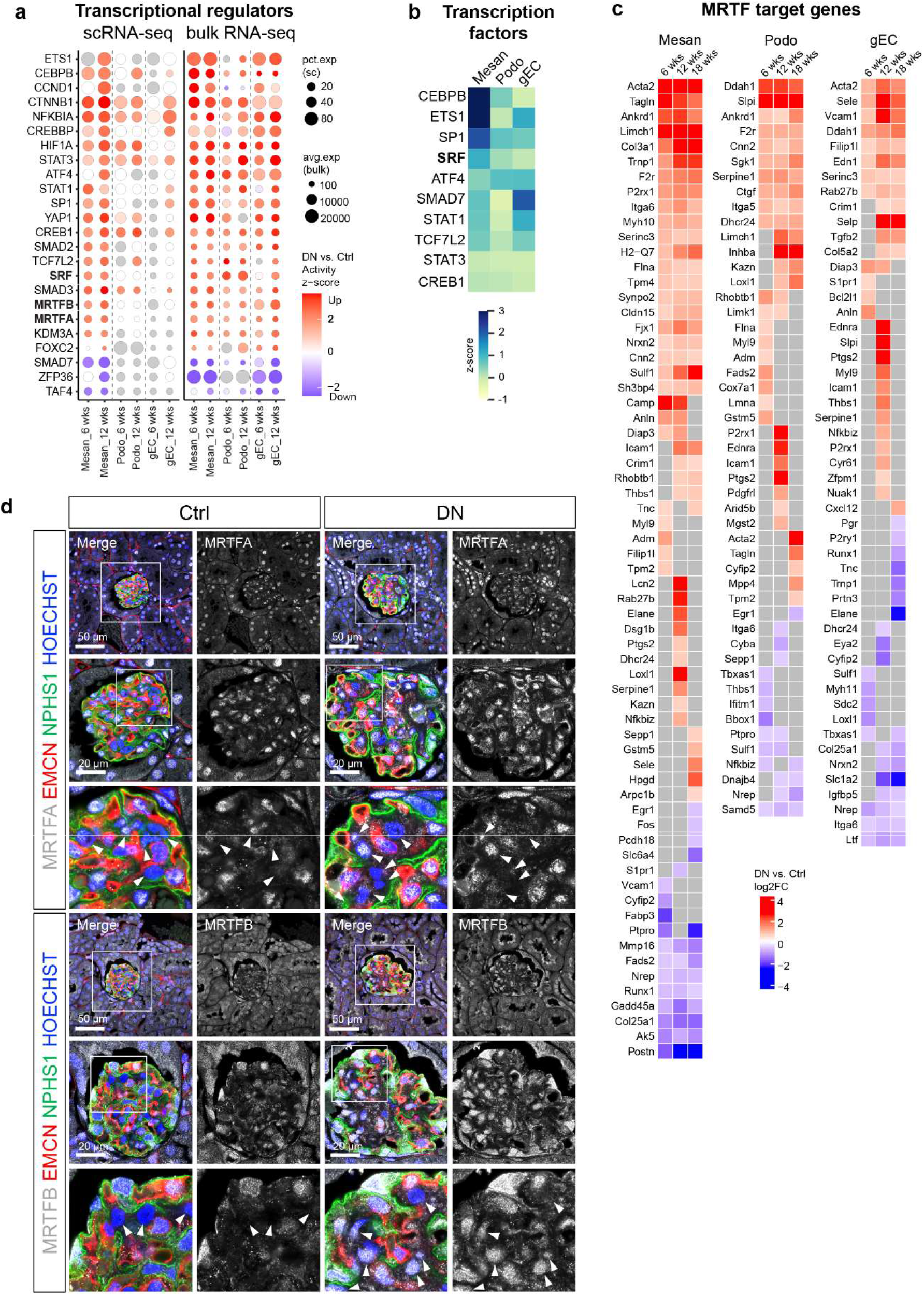
Mechanosensitive transcriptional regulators are activated in DN glomeruli. (a) Dot plot displaying the commonly changed transcriptional regulations in both single-cell and bulk RNA-seq data estimated by IPA. (**b**) Heatmap showing the activity z scores of transcription factors estimated by SCENIC. (**c**) Heatmap showing the log2FC (DN vs. Ctrl) values of MRTF transcriptional target genes in mice glomerular cells. Nonsignificant genes are shown in grey (**a, c**). (**d**) Immunofluorescence staining of the mouse kidney paraffin sections showing the expression and localization of MRTFA and MRTFB in control and DN glomeruli. Glomerular endothelial cells were labelled by EMCN (endomucin), podocytes were labelled by NPHS1 (Nephrin), nuclei were counterstained with HOECHST. Arrows indicate mesangial cells, which were negative for EMCN and NPHS1.

Additionally, the gene regulatory network was inferred using SCENIC [18, 19] to determine the cell-type specific activity of transcription factors (Supplementary Fig. 6b). Consistently, the activity of SRF was high in mesangial cells, and moderate or low in podocytes and gECs, respectively (Fig. 3b).

The MRTF-SRF transcriptional target genes with significant regulations in DN glomerular cell types are shown in Figure 3c. Notably, the MRTF-SRF-specific targets *Acta2* and *Tagln* [20] were found in both single-cell and bulk RNA-seq data. Enrichment analysis of significantly changed MRTF transcriptional target genes suggested regulations of smooth muscle proliferation, actin cytoskeleton, integrin cell surface interactions and extracellular matrix organization etc. (Supplementary Fig. 6c).

MRTF-SRF is an important mechanosensitive transcriptional pathway in cells [21]. MRTF family consists of myocardin, MRTFA (MKL1, MAL) and MRTFB (MKL2). Unlike myocardin, which is limited to myocardial cells and smooth muscle cells, MRTFA and MRTFB are found in various cells and tissues including the kidney [22]. Mechanical stresses lead to the translocation of cytoplasmic MRTF into the nucleus to initiate the SRF-mediated transcriptional responses [22]. We investigated the expression and location of MRTFA and MRTFB in mouse DN by immunofluorescent staining. MRTFA and MRTFB were found in the cytoplasm and nuclei of podocytes and gECs, exhibiting comparable expression levels in both control and DN mice (Fig. 3d). Mesangial cells exhibited a low expression level of MRTFA in control mice, while some of the mesangial cells exhibited nuclear accumulation of MRTFA in DN mice (Fig. 3d). MRTFB expression was absent in most mesangial cells in control mice, but its expression level was clearly increased in DN mice. Importantly, nuclear accumulation of MRTFB in mesangial cells was obvious to detect especially in the region where mesangial expansion was taking place (Fig. 3d).

### MRTF transcriptional target genes are upregulated in DN patients with type 2 diabetes

We investigated the expression of MRTF transcriptional target genes in an early DN cohort of Pima Indians from the Gila River Indian Community in Arizona, individuals with type 2 diabetes [23]. mRNA expression analysis of microdissected glomeruli showed that various genes significantly changed in mouse DN were also significantly regulated in early human DN (Fig. 4a). Many of these target genes were positively correlated with mesangial volume (Fig. 4b). Independent single-nucleus transcriptomics data from early human DN [3] showed that these MRTF transcription genes were mostly regulated in mesangial cells (Supplementary Fig. 7a). The MRTF transcriptional target genes were mapped in the ERCB dataset. These genes were significantly regulated in glomeruli in the context of DN and other diseases potentially causing mechanical stresses, such as arterial hypertension, as well as various other glomerular diseases with glomerulosclerosis, such as lupus nephritis (caused by systemic lupus erythematosus, SLE) and IgA nephropathy, but not in minimal change disease (MCD) (Fig. 4c).

**Fig. 4:**
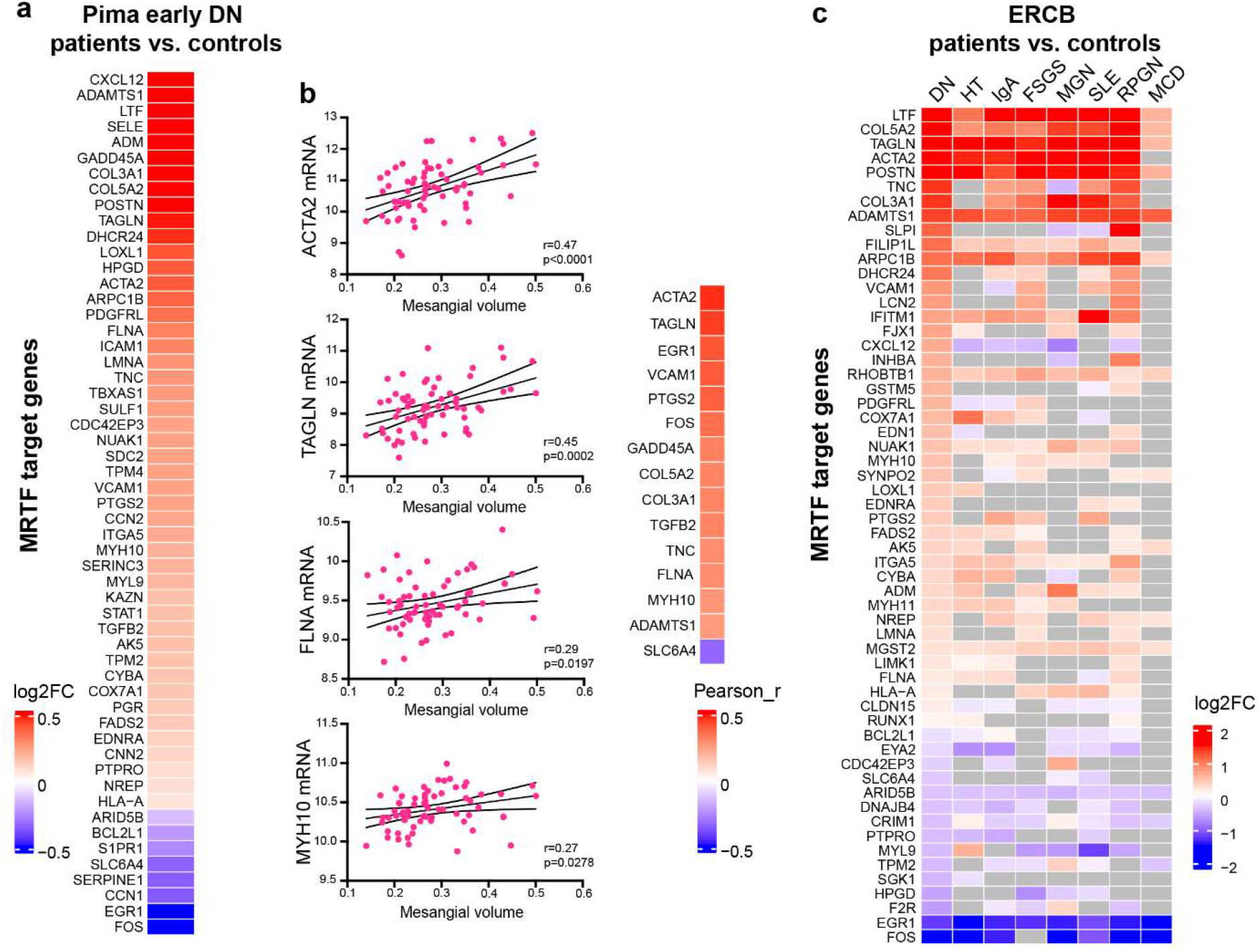
MRTF transcriptional target genes are upregulated in DN patients with type 2 diabetes. (**a**) Heatmap showing the log2FC (DN vs. Ctrl) values of MRTF transcriptional target genes in microdissected glomeruli from Pima early DN patients. (**b**) Pearson correlations between MRTF target gene expression levels and mesangial volumes in Pima early DN patients (n= 66). (**c**) Heatmap showing the log2FC (DN vs. Ctrl) values of MRTF transcriptional target genes in microdissected glomeruli from ERCB patient datasets. Nonsignificant genes are shown in gray. DN: diabetic nephropathy, HT: hypertensive nephropathy, IgA: IgA nephropathy, FSGS: focal segmental glomerulosclerosis, MGN: membranous nephropathy, SLE: lupus nephritis, RPGN: ANCA-associated glomerulonephritis, MCD: minimal change disease.

Next, we investigated the expression and location of MRTFA and MRTFB in early human DN by immunofluorescent staining. A basal and comparable expression level of MRTFA was detected in all three glomerular cell types in both healthy and early DN kidney biopsy samples (Fig. 5). Consistent with the observations in mouse DN, MRTFB was absent in most mesangial cells from healthy individuals but was accumulated in the nuclei of mesangial cells where glomerular lesions were observed (Fig. 5). Taken together, our immunofluorescent staining analyses on human and mouse DN confirm the activation of MRTFB in mesangial cells during early DN.

**Fig. 5:**
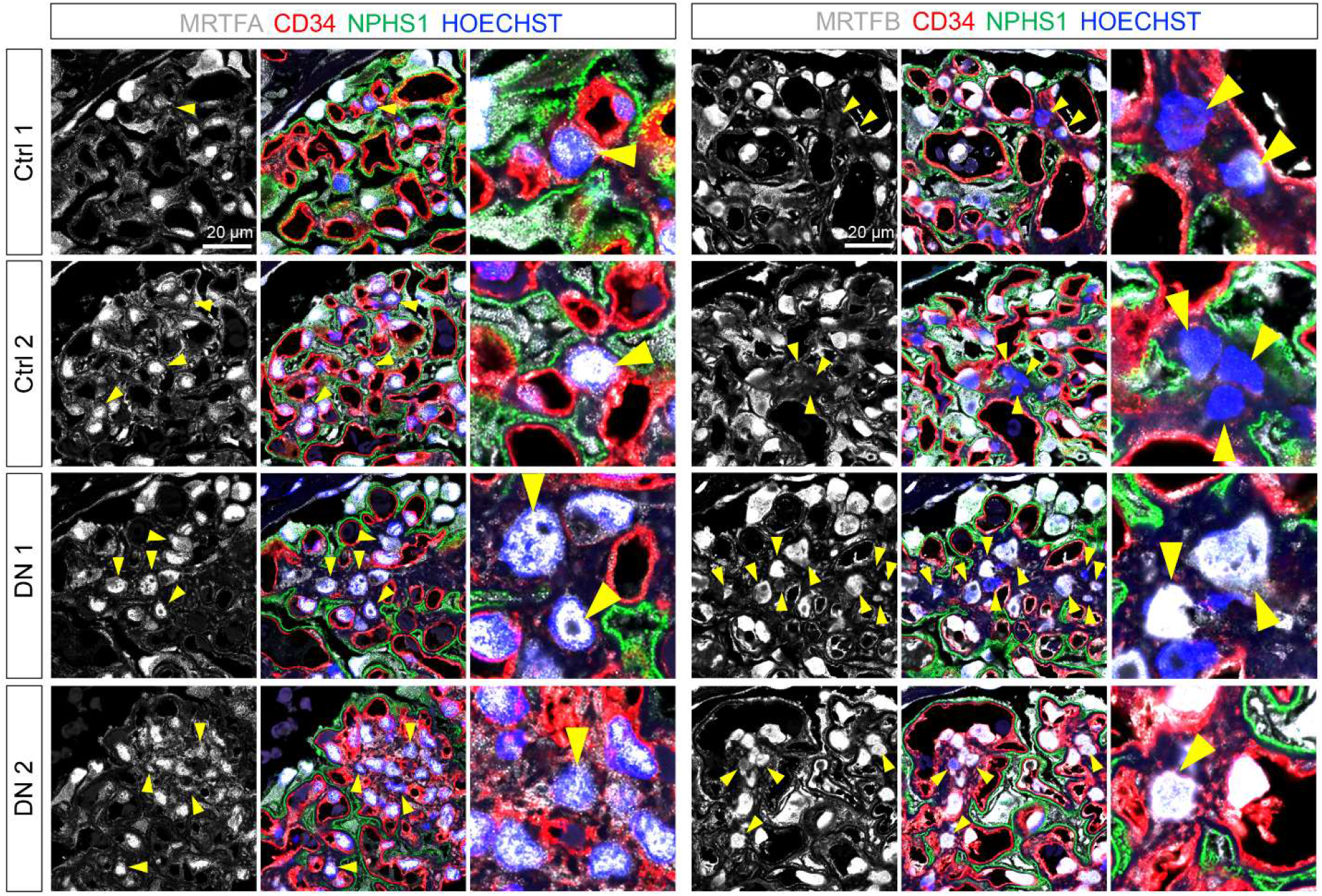
Immunofluorescence staining confirms the activation of MRTFB in mesangial cells during early DN. Immunofluorescence staining of the human kidney samples (ctrl: n=2, DN: n=5). Representative images showing the expression and localization of MRTFA and MRTFB in control and DN glomeruli. Glomerular endothelial cells were labelled by CD34, podocytes were labelled by NPHS1 (Nephrin), nuclei were counterstained with HOECHST. Arrows indicate mesangial cells, which were negative for CD34 and NPHS1.

### Mesangial cells exhibit dominant signaling networks in DN glomeruli

Our results suggested that MRTF-SRF transcriptional regulation was activated in both mouse and human DN, predominantly in mesangial cells. Additionally, we identified various mesangial cell marker genes that encode transmembrane proteins responsible for the perception and transduction of mechanical signals, including mechanosensitive ion channels (MSCs), GPCRs, integrins and cadherins [21] (Fig. 6a). Furthermore, we performed CellChat [24] to study the changes of intercellular communications (secreted signaling, cell-cell contact and ECM-receptor) taking place in DN. We compared interaction strength between all kidney cell types in DN versus control mice, which showed dominating interactions of mesangial cells with all other cell types in DN kidneys (Fig. 6b). Additionally, mesangial cells dominated the intercellular communications (Fig. 6c) and interacted as a sender extensively with gECs and podocytes (Fig. 6d) in DN glomeruli. These interactions involved various well-known growth factor signaling pathways associated with the pathogenesis of chronic kidney disease (CKD), such as bone morphogenetic protein (BMP), fibroblast growth factor (FGF) and vascular endothelial growth factor (VEGF), as well as interactions between multiple extracellular matrix proteins and integrins. Notably, the BMP2/4 signaling pathways identified in DN mesangial cell interactions contribute to glomerulosclerosis and tubulointerstitial fibrosis, particularly in DN [25].

**Fig. 6:**
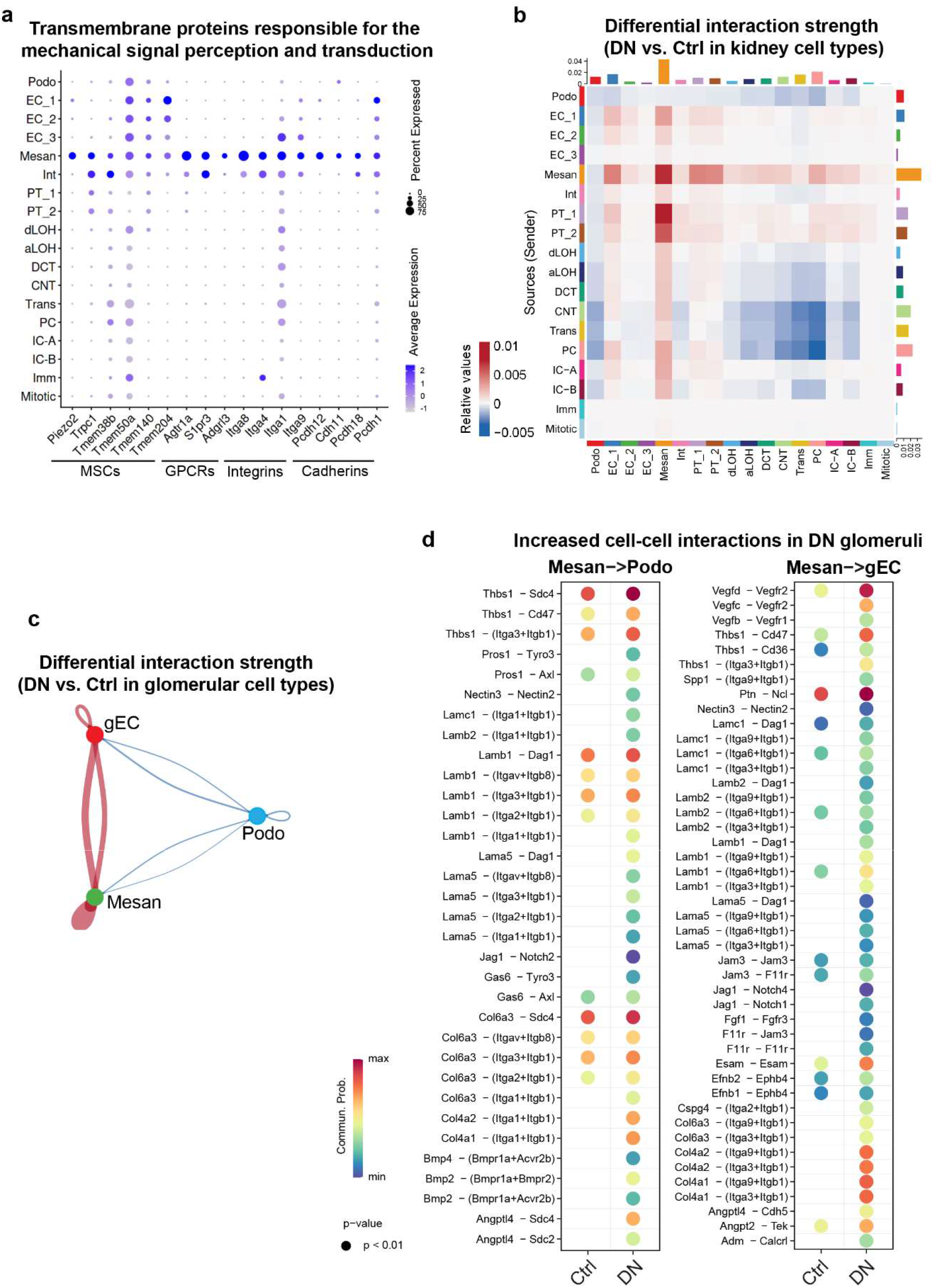
Mesangial cells exhibit dominant signaling networks in DN glomeruli. (**a**) Dot plot showing mesangial marker genes encoding transmembrane proteins responsible for the perception and transduction of mechanical signals. MSCs: mechanosensitive ion channels, GPCRs: G-protein coupled receptors. (**b**) Change of cell-cell interactions in all pairs of kidney cell types in DN vs. Ctrl mice. (**c**) Change of cell-cell interactions in all pairs of glomerular cell types in DN vs. Ctrl mice. (**d**) Increased cell-cell interactions for mesangial cell signaling to podocytes and glomerular endothelial cell in DN mice.

### Kidney ex vivo perfusion activates mechanosensitive signaling pathways

Glomerular hyperfiltration as a consequence of increased single-nephron glomerular filtration rate may be a common upstream mechanism contributing to CKD [2]. In diabetes, glomerular hyperfiltration is driven by the sodium–glucose cotransporter 2 (SGLT2) [2]. Hyper-reabsorption of glucose and sodium in the PT ultimately leads to restricted tubuloglomerular feedback, resulting in increased intraglomerular pressure and glomerular hyperfiltration [2]. Consequently, circumferential and axial capillary wall stress, as well as fluid shear stress at the glomerular filtration barrier is increased [26]. Since the mechanosensitive signaling pathway MRTF-SRF is a central coordinator of fibrosis-relevant mechanoresponse [21], we hypothesize that glomerular hyperfiltration may directly activate MRTF-SRF, leading to cellular changes associated with diabetic glomerulopathy.

We used normothermic machine perfusion to proof this hypothesis. The ex vivo kidney perfusion takes place for 1 to 2 hours at 100 mmHg with 100% Gey’s Balanced Salt Solution (GBSS) without the addition of albumin or red blood cells (Methods). Under this condition, tubular glomerular feedback is absent and 100 mmHg is a supraphysiologic pressure leading to glomerular hyperfiltration [27].

We generated bulk and single-nucleus transcriptomics data of isolated mouse glomeruli and pig renal cortex tissue, respectively (Fig. 7a, d), and gene expression was compared in perfused versus unperfused conditions. In ex vivo perfused mouse glomeruli, upstream analysis in DEGs suggested activation of MRTFA/B, SRF and YAP1 (Fig. 7b). Enrichment analysis of significantly changed MRTF transcriptional target genes revealed actin cytoskeleton organization, VEGFA-VEGFR2 signaling pathway, smooth muscle contraction, PDGFRB pathway and cellular response to external stimulus etc. (Fig. 7c). In ex vivo perfused pig kidney, a total of 11,692 single nuclei were profiled from perfused and unperfused renal cortex tissue. Unsupervised clustering identified 18 clusters (Supplementary Fig. 7b), which were classified into 12 cell types (Fig. 7e; Supplementary Fig. 7c) on the basis of cell/nucleus-specific marker genes reported for previous human kidney single-nucleus data (Fig. 7f) [3, 16]. We identified the major glomerular cell types including a stromal cell type (indicated by “STROMA” in figures) defined by the nucleus marker gene MEIS1 and FHL2 (Fig. 7f) [28]. Immunofluorescent staining on the pig renal cortex showed that MEIS1-positive cells were mainly composed of mesangial cells, but also some interstitial cells (Fig. 7g). Upstream analysis on DEGs revealed the activation of MRTFA/B, SRF and YAP1 in stromal cells, as well as the activation of YAP1 and MRTFB in podocytes and endothelial cells, respectively (Fig. 7h). Significantly regulated MRTF transcriptional target genes were found predominantly in stromal cells (Fig. 7i). These findings support our hypothesis that the activation of mechanosensitive signaling pathways is a common mechanism in response to glomerular hyperfiltration.

**Fig. 7:**
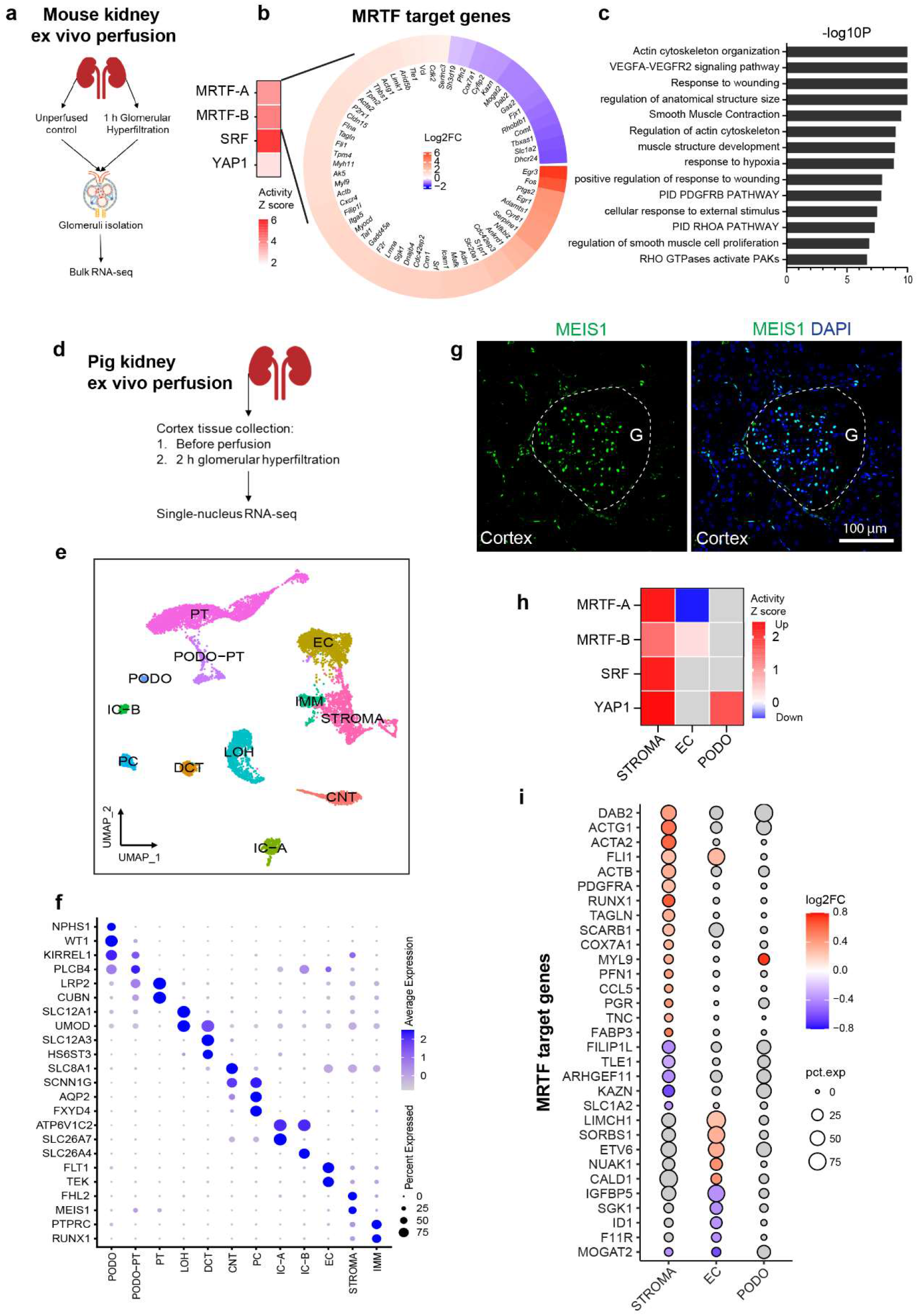
Kidney ex vivo perfusion activates mechanosensitive signaling pathways. (**a**) Experimental scheme of mouse kidney ex vivo perfusion. (**b**) Heatmap showing the log2FC values of MRTF transcriptional target genes in perfused versus control glomeruli. (**c**) Top enriched pathways of MRTF transcriptional target genes. (**d**) Experimental scheme of pig kidney ex vivo perfusion. (**e**) UMAP plot of annotated cell types from snRNA-seq of perfused and control pig kidney tissue. (**g**) Immunofluorescence staining against MEIS1 in the pig kidney cortex. Nuclei were counterstained with DAPI. G: Glomerulus. Dashed lines indicate the area of glomerulus. Scale bar: 100 μm. (**f**) Dot plot displaying defining marker genes for each cell type. STROMA: stromal cells. (**h**) Heatmap showing z scores of mechanosensitive transcriptional regulators estimated by IPA. (**i**) Dot plot displaying MRTF transcriptional target genes significantly changed in perfused kidney tissue. Nonsignificant regulation or genes are shown in gray (**g, h**).

## Discussion

BTBR *ob/ob* is a useful mouse model to investigate cellular changes and molecular mechanisms in early DN. Integrative analysis of single-cell and bulk RNA-seq data led to the identification of the mechanosensitive pathway MRTF-SRF in DN glomeruli. The association of MRTF-SRF activation with human DN was validated in independent patient cohorts at both mRNA and protein levels, and MRTF-SRF is suggested as a common mechanism in response to glomerular hyperfiltration.

Single-cell analysis on BTBR *ob/ob* kidneys suggested that PT cells and mesangial cells are the most sensitive cells to hyperglycemia and initiate glucose-dependent and glucose-independent downstream pathways in response to stress in the kidneys, respectively. PT cells showed adaptations to glucose metabolic fluxes and upregulation of antioxidant pathways. Increased glucose influx in cells fuels oxidative phosphorylation, leading to overproduction of superoxide by mitochondria and generation of reactive oxygen species, which play a central role in initiating diverse pathways responsible for diabetic abnormalities such as cellular dysfunction, inflammation and fibrosis [1]. Mesangial cells showed the activation of other signaling pathways obviously independent of glucose metabolic flux. This finding is consistent with those of previous studies showing that overexpression of GLUT 1 in mesangial cells exposed to high glucose fails to induce TGF-β1 synthesis and oxidative stress [29] and that transgenic mice with GLUT1 overexpression in mesangial cells do not develop pathological phenotypes until 26 weeks of age [30].

Mouse DN glomeruli exhibited shared features with human DN glomeruli. Approximately 70% glomerular DEGs identified in DN mice were also significantly regulated in DN patients. Importantly, MRTF-SRF transcriptional target genes were significantly regulated in both mouse and human DN glomeruli. Notably, mesangial cells responded at the earliest time point and showed a persistent activation of MRTFB compared to podocytes and gECs. Immunofluorescent staining further confirmed the activation of MRTFB in the region where mesangial expansion was taking place. Mechanic stimulation triggers cytoskeleton remodeling, resulting in the activation of mechanosensitive transcriptional pathway MRTF-SRF [21], which plays a key role in organ fibrosis [21, 31]. Upon stimulation, cytoplasmic MRTF translocate to the nucleus and interact with SRF to target the expression of muscle-specific and contractile genes, as well as extracellular matrix genes [32, 33]. Interestingly, single-cell data showed mesangial cell expression of numerous genes that encode transmembrane proteins responsible for the perception and transduction of mechanical signals. DN mesangial cells exhibited dominant signaling network in the whole kidney, interacted with gECs and podocytes by ECM and secreted signaling. These results suggest that MRTF-SRF transcriptional regulation in mesangial cells could be an important pathological pathway contributing to diabetic glomerulopathy.

Since the mechanical stress caused by glomerular hyperfiltration is thought to play a critical role in the development of glomerulopathy [26], we speculate that glomerular hyperfiltration, a key event in early DN [1, 2], directly activates mechanosensitive signaling pathways in renal glomeruli, and consequently contribute to diabetic glomerulopathy. This speculation is supported by evidence in our study. First, single-cell and bulk RNA-seq of DN mice showed activation of MRTFA/B in three major glomerular cell types, which was also observed in an early DN cohort. Second, ex vivo kidney perfusion induced glomerular hyperfiltration showed a prominent activation of MRTFA/B, SRF and YAP1 in mouse glomeruli. Third, snRNA-seq of perfused pig renal cortex tissue showed activation of these mechanosensitive transcriptional regulators, which were most prominently activated in stromal cells composing mesangial cells.

Many MRTF target genes were significantly regulated not only in DN but also in the context of other glomerular diseases associated with glomerulosclerosis. These findings suggest a common role of MRTF in diabetic glomerulopathy, as well as in glomerulosclerosis generally. Previous studies demonstrated that MRTF-SRF and YAP are essential for podocyte structure and function [34, 35]. Tubulointerstitial fibrosis was diminished in MRTFA-deficient mice and MRTFA was activated in high-fat diet- and streptozotocin-induced DN [36]. In addition, genetic deletion or pharmacological inhibition of MRTF attenuates fibrosis in various pathological conditions, including kidney diseases like diabetic nephropathy, obstructive nephropathy, acute kidney injury, and polycystic kidney disease [31]. Therefore, MRTF is important for glomerular cells and is a potential therapeutic target for DN, as well as glomerulosclerosis.

Finally, while the principle of SGLT-2 inhibition has been proven to be the major advance in slowing down CKD, the mechanisms remain unclear [2]. It was proposed that a lowering of the intraglomerular pressure by SGLT-2 inhibition might drive the renal protective effects. Our results unravel now mechanosensitive signalling pathways as possible mediators of SGLT-2 dependent effects, opening the search of novel druggable downstream targets of SGLT-2.

In conclusion, our study presents a comprehensive single-cell transcriptomic landscape of early DN and reveals cell-specific alterations in gene expression that occur with DN onset and progression. Our intensive analysis of glomerular cell types reveals that the mechanosensitive signaling is associated with diabetic glomerulopathy and could play a driving role in response to glomerular hyperfiltration. The MRTF transcription pathway may be part of a common mechanism in the glomerulus in the context of glomerulosclerosis.

## Methods

### Animals

All animal experiments were conducted according to the National Institutes of Health Guide for the Care and Use of Laboratory Animals as well as the German law for the welfare of animals. Mice were housed in a specific pathogen-free (SPF) facility with free access to chow and water and a 12 h/12 h day/night cycle. Breeding and genotyping were performed according to standard procedures.

#### BTBR ob/ob podocyte-reporter mice

Animal experiments were approved by local authorities (Ü003-2018/BGV Hamburg). BTBR *ob/+* (BTBR.Cg-Lep^ob/wt WiscJ^) heterozygous animals (Jax No. 004824) and corresponding wild-type mice BTBR (Jax No 002282) were purchased from JAX (Bar Harbor, Maine, USA). Podocyte-reporter mice (Gt(ROSA)26Sor^tm4(ACTB-tdTomato,-EGFP)Luo^;Tg(NPHS2-cre)^295Lbh^) were also purchased from JAX (Jax No. 007576) and crossed for at least 7 generations with BTBR wild-type animals before BTBR.Cg-Lep^ob/wt WiscJ^;Gt(ROSA)26Sor^tm4(ACTB-tdTomato,-EGFP)Luo^;Tg(NPHS2-cre)^295Lbh^ animals were generated. The resultant offspring was intercrossed to yield BTBR.Cg-Lep^ob/ob WiacJ^;Gt(ROSA)26Sor^tm4(ACTB-tdTomato,-EGFP)Luo^;Tg(NPHS2-cre)^295Lbh^ and BTBR.Cg-Lep^wt/wt WiscJ^;Gt(ROSA)26Sor^tm4(ACTB-tdTomato,-EGFP)Luo^;Tg(NPHS2-cre)^295Lbh^ animals.

#### Mouse kidney normothermic machine perfusion

All procedures involving mouse kidney perfusion described in this manuscript were conducted according to German- and Hamburg law and approved by the veterinary administration of the City of Hamburg under the license N002/2020. As previously described [37], male C57BL/6 kidneys were collected under anesthesia (i.p. injection of 100 μL/10g body weight of a solution containing 10 mg/ml Ketamine and 1,6 mg/ml Xylazine). Briefly, a laparotomy was performed, the aorta was cannulated, ligatures placed and closed around all main vessels and perfusion initiated only in the right kidney. The left unperfused kidney was then removed and used as control. After this, the right kidney was removed from the mouse during ongoing perfusion. Perfusion took place ex vivo at 100 mmHg at 37°c with 100% GBSS solution without the addition of albumin or red blood cells (Sigma G9779) continuously gassed in a dialyzer with 100% carbogen (95% O2, 5% CO2). Perfusion was stopped after 60 minutes. Isolation of glomeruli was performed as described in supplementary methods (FACS-sorted glomerular cells).

#### Pig kidney normothermic machine perfusion

Pig kidneys were collected from a German slaughterhouse. Kidneys were received directly after slaughter and immediately flushed with 400-500 ml 1x PBS with 2 ml Heparin 10.000 IE/ml. Kidney were stored at +4°c for around 2 hours, after which normothermic perfusion was initiated. Perfusion took place at 100 mmHg at 37°c with increased glomerular pressure and hyperfiltration as described above. Renal cortex tissue was collected either before perfusion (control) or 120 minutes after start of perfusion (perfused).

### Single-cell data analysis

#### Preprocessing and quality control (QC) of scRNA-seq data

10x Genomics raw sequencing data were processed using CellRanger software (version 3.0.2, 10x Genomics, Pleasanton, CA), and the 10x Genomics mouse genome mm10 3.0.0 release was used as the reference genome (function cellranger count). The matrices of cells and the unique molecular identifier (UMI) count were obtained and further processed with the R package Seurat (version 3.1.1) [39]. For QC, we first filtered out genes detected in fewer than 3 cells and data for cells in which fewer than 200 genes had nonzero counts. To remove potential doublets, cells with more than 7,000 expressed genes (nFeature) were excluded. We removed low-quality cells with more than 50% mitochondrial genes among all detected genes, as is recommended for kidney tubular cells [40]. After clustering and cell type identification, we performed curated doublet and high-mitochondrial gene cell removal (see below) based on known lineage-restricted markers.

#### Dimensionality reduction and clustering

The Seurat R package (version 4.0.2) was used to perform unsupervised clustering analysis on scRNA-seq data. In brief, gene counts for cells that passed QC were normalized by library size and log-transformed (function NormalizeData, normalization.method = “LogNormalize”, scale.factor = 10,000). Then, highly variable genes were detected (function FindVariableFeatures, selection.method = “vst”, nfeatures = 2,000). To reduce batch effects, we applied the integration method implemented in Seurat version 3 (functions FindIntegrationAnchors and IntegrateData, dims = 1:30). The integrated matrix was then scaled with the ScaleData function (default parameters). PCA was performed on the scaled data (function RunPCA, npcs = 30) to reduce dimensionality.

The number of principal components used for each clustering round was dataset dependent and was determined on the basis of the elbow of a PCA scree plot. The selected principal components were then used to compute the KNN graph based on the Euclidean distance (function FindNeighbors). Cell clusters were subsequently generated using the function FindClusters. The resolution of the FindClusters function for each dataset was also determined by exploration of the top marker genes of each cluster. Uniform Manifold Approximation and Projection (UMAP) was used to visualize the clustering results. The top DEGs in each cluster were found using the FindAllMarkers function (min.pct = 0.25, logfc.threshold = 0.25) with Wilcoxon rank-sum tests. The most highly expressed genes were then used to determine the cell type of each cluster.

#### Curated doublet and high-mitochondrial gene cell removal

After the cell type was determined for clusters, we performed additional curated doublet and high-mitochondrial-cell removal. Based on the literature and the exploration of our datasets, we created a lineage-restricted marker gene list for tubular (*Cubn, Epcam*) and nontubular (*Pecam1, Pdgfrb, Nphs2, Ptprc*) cell types. We removed the cells that expressed markers of the opposite lineage. For nontubular cells, we discarded cells with more than 20% mitochondrial genes among all detected genes. There were 86,508 cells before curated doublet removal, 71,831 cells after curated doublet removal and 770,944 cells after high-mitochondrial gene cell removal.

#### Differential gene expression analysis

The Seurat FindMarker function was used to perform differential gene expression analysis for each cell type between the control and DN groups. The test method used in the FindMarker function was MAST (v1.12.0) [41].

### Single-nucleus data analysis

#### Preprocessing and QC

10x Genomics raw sequencing data were processed using CellRanger software (version 5.0.1, 10x Genomics, Pleasanton, CA). Pig genome was built by function “cellranger mkref” using the Sscrofa v103 FASTA file and GTF file (https://www.ensembl.org) and then mapping was done and the count matrices were generated by function “cellranger count” with parameter “include-introns”. For QC, firstly we applied soupX [42] with default parameters to remove ambient RNA contamination. Then the nuclei were filtered out if the number of genes detected was less than 500 or greater than 6000 or the percentage of mitochondrial genes detected exceeded 7%. To further remove potential doublets, Scrublet [43] was applied to the data and cells were excluded if they were identified as doublets (default parameters).

#### Dimensionality reduction and clustering

An updated version of Seurat R package (version 4.0.2) was used to perform unsupervised clustering analysis on snRNA-seq data since the experiments were performed later than the mouse scRNA-seq analysis. Count data normalization, scaling, highly variable genes selection and sample integration were same as the mouse single cell data process described above. PCA was performed on the scaled data (function RunPCA, npcs = 30) and the first 15 PCs were used for clustering. The KNN graph was calculated (function FindNeighbors) and then the clustering result is obtained (function FindClusters, resolution = 0.5). To visualize the clustering result, UMAP coordinates were calculated (function RunUMAP, dims = 1:15). Wilcoxon method was used to perform the differential gene analysis. For each cluster, the marker gene list was determined by log2FoldChange > 0.25 and adj.p value < 0.05 (FindAllMarkers). The most highly expressed genes were then used to determine the cell type of each cluster.

### Bulk RNA-seq

#### Library preparation and RNA-seq

A small amount of RNA (2 ng) was used as input material, and libraries were prepared with a SMART-Seq Stranded Kit according to the user manual (Takara Bio USA, Mountain View, CA, USA). In brief, samples were fragmented at 85 °C for 6 min prior to first-strand synthesis. Illumina adaptors and indexes were added to single-stranded cDNA via 5 cycles of PCR. After library purification with AMPure beads and depletion of ribosomal cDNA with scZapR, final RNA-seq library amplification (13 cycles) was conducted, and the final RNA-seq library was purified with AMPure beads. The library samples were quantified using Quant-iT PicoGreen dsDNA Reagent (Invitrogen; Thermo Fisher Scientific, Waltham, MA, USA) on a ClarioStar microplate reader according to the manufacturer’s instructions (BMG LABTECH, Ortenberg, Germany). The quality, including fragment size, of the cDNA was assessed on an Agilent Technologies Bioanalyzer 2100 using an Agilent DNA 1000 kit according to the manufacturer’s instructions (Agilent Technologies, Palo Alto, CA, USA).

Pooled samples were quantified with a Qubit 1X dsDNA HS Assay Kit on a Qubit fluorometer (Thermo Fisher Scientific, Waltham, MA, USA). Single-read sequencing was performed on a NovaSeq 6000 device using an S2 Reagent kit (100 cycles) according to the manufacturer’s instructions (Illumina Inc., CA, USA).

#### Bulk RNA-seq data analysis

The quality of the bulk RNA-seq reads was assessed using FastQC (v0.11.5), and the reads were aligned to the mouse reference genome (mm10) with Bowtie2 (v2.3.3.1) [44] using RSEM (v1.3.0) [45] with the default parameters. The function rsem-calculate-expression was used to align the reads and quantify the gene and isoform abundance. The output of rsem-calculate-expression separately gives the read count and transcripts per million (TPM) value for each gene and isoform. Differential expression analysis was carried out using gene read counts with the DESeq2 package (v1.22.2) [46] to produce log2FC values and corresponding p values and adjusted p values. PCA was performed using regularized log transformation of the count data, and the results were visualized using gplots (v3.0.1.1).

### Integrated analysis

#### Correlation of bulk RNA-seq and scRNA-seq data

For podocytes, ECs and mesangial cells, the single-cell data were correlated with bulk RNA-seq data. The cell type markers (top DEGs of each cell type identified by the Seurat FindAllMarkers function (min.pct = 0.25, logfc.threshold = 0.25) were used for correlation analysis. The normalized gene expression count matrices of cell type markers were extracted from the single-cell data and bulk RNA-seq data separately, and then Pearson correlation coefficients were calculated between every pair of single cells and replicates of bulk RNA-seq data.

#### CellChat

We applied Cellchat (v 1.4.0) [24] to infer cell-cell communications across all kidney cell types and the glomerular cell types. We used the mouse database curated in CellChat including the “Secreted Signaling”, “Cell-cell Contact” and “ECM-Receptor”. The comparison analysis between diseased samples and control samples was performed according to the Cellchat tutorial (https://htmlpreview.github.io/?https://github.com/sqjin/CellChat/blob/master/tutorial/Comparison_analysis_of_multiple_datasets.html). The communication probabilities were compared to identify the unregulated and downregulated signaling ligand-receptor pairs between the two conditions.

#### SCENIC

The gene regulatory network was inferred using pySCENIC (v 0.11.2, a lightning-fast python implementation of the SCENIC pipeline [18, 19]. Firstly, the GRNboost2 algorithm was used to infer gene regulatory network and generate coexpression modules (pyscenic grn). Next, the regulon prediction step was performed (pyscenic ctx, using the default parameters and mm10__refseq-r80 10kb_up_and_down_tss.mc9nr and mm10 refseq-r80__500bp_up_and_100bp_down_tss.mc9nr motif collections). Finally, the AUCell matrix was generated with a threshold of 0.01 (pyscenic aucell). Cell type specific regulators were identified based on Z-score of the AUCell values for the cells of a give type.

#### Upstream analysis

We performed Qiagen’s IPA tool to identify transcriptional regulators (significance: P value <0.05). Positive and negative z-score values indicated the activation or inhibition of transcriptional regulators, respectively.

#### Enrichment analysis

GO biological processes, KEGG, WikiPathways, and Reactome Gene Sets were performed with Metascape [47].

### Human data

#### Microarray analysis

Human renal indication biopsy specimens were collected in an international multicenter study and deposited in the ERCB-Kröner-Fresenius Biopsy Bank (ERCB-KFB [48]; for participating centers, see Shved *et al* [49]). Protocol kidney biopsy specimens were collected from Pima Indians with type 2 diabetes. The study subjects participated in an intervention trial (Renoprotection in Early Diabetic Nephropathy in Pima Indians, clinicaltrials.gov, NCT00340678) [23, 50]. Biopsies were obtained from patients after informed consent was obtained and with approval of the local ethics committees. Tissue processing and microdissection protocols were performed similarly on both ERCB and Pima biopsy material. Following renal biopsy, the tissue was transferred to an RNase inhibitor and microdissected into glomeruli and tubulointerstitial tissue. Total RNA was isolated, reverse-transcribed and amplified. Fragmentation, hybridization, staining and imaging were performed according to the Affymetrix Expression Analysis Technical Manual (Affymetrix, Santa Clara, CA, USA). Glomerular samples from patient with different renal diseases were analyzed for mRNA expression levels (GSE32591, GSE35489, GSE37463, GSE47185, GSE99340). The analysis included gene expression profiles from patients with DN (n=14 glomeruli), hypertensive nephropathy (HT; n=15 glomeruli), MCD (n=14 glomeruli), IgA nephropathy (IgA; n=27 glomeruli), FSGS (n=23 glomeruli), membranous nephropathy (MGN; n=21 glomeruli), lupus nephritis (SLE; n=32 glomeruli), ANCA-associated glomerulonephritis (RPGN; n=23 glomeruli), and controls (LDs; n=41 glomeruli). To account for ethical considerations, to ensure privacy protection, and to avoid identifying individual study participants in this vulnerable population of Pima people, the Institutional Review Board of the National Institute of Diabetes and Digestive and Kidney Diseases has stipulated that individual-level gene expression and genotype data from this study cannot be made publicly available. CEL file normalization was performed with the robust multichip average method using RMAExpress (Version 1.0.5) and the human Entrez-Gene custom CDF annotation from Brain Array version 18 (PIMA) and 25 (ERCB) (http://brainarray.mbni.med.umich.edu/Brainarray/default.asp). The log-transformed ERCB dataset was corrected for batch effect using ComBat from the GenePattern pipeline (http://www.broadinstitute.org/cancer/software/genepattern/). To identify DEGs, the Significance Analysis of Microarrays (SAM) method [51] was applied using the SAM function in Multiple Experiment Viewer (TiGR MeV, Version 4.9). A q-value below 5% was considered to indicate statistical significance.

## Acknowledgments

The authors wish to thank Alina Borchers, Anja Obser, Silvia Chilla and Valerie Oberueber, as well as FACS Sorting Core Unit and Microscopy Imaging Facility at University Medical Center Hamburg-Eppendorf, Germany, for the excellent technical support. TBH was supported by the DFG (CRC1192, HU 1016/8-2, HU 1016/11-1, HU 1016/ 12-1), by the BMBF (STOP-FSGS-01GM1901C, NephrESA-031L0191E and DEFEAT PANDEMIcs), by the Else-Kröner Fresenius Foundation (Else Kröner-Promotionskolleg –iPRIME), and by the European Research Council-ERC (grant 616891). This work was further supported by Unicyte AG and by the H2020-IMI2 consortium BEAt-DKD (115974); this joint undertaking receives support from the European Union’s Horizon 2020 research and innovation program and EFPIA and JDRF (TBH). The work performed at the University of Michigan was supported by George M. O’Brien Michigan Kidney Translational Core Center funded by NIH/NIDDK grant 2P30-DK-081943. The work performed in Phoenix was supported by the Intramural Research Program of the National Institute of Diabetes and Digestive and Kidney Disease and the American Diabetes Association (Clinical Science Award 1-08-CR-42). The ERCB-KFB was supported by the Else Kröner-Fresenius Foundation. We also thank all participating centres of the European Renal cDNA Bank – Kröner-Fresenius biopsy bank (ERCB-KFB) and their patients for their cooperation. Active members at the time of the study are found in N. Shved et al., Scientific reports 7, 8576 (2017). This work was partially funded by the Else Kröner-Fresenius-Stiftung and the Eva Luise und Horst Köhler Stiftung - Project No: 2019_KollegSE.04.

## Author contributions

Conceptualization: SL and TBH. Formal analysis: SL, SLU and YZ. Investigation: SL, SLU, YZ, MTL, VN, JC, TZ, HA, MS, SZ, SEG, GW, ZL, TW and DD. Method development: SL, SLU, CMS and CK. Resources: MTL, VN, RGN, DF, TW, FG and MK. Data and sample curation: SL, SLU, YZ, MTL, VN and SB. Writing (original draft): SL and TBH. Visualization: SL. Supervision and funding acquisition: TBH.

## Notes

### Competing Interest Statement

The authors have declared no competing interest.

